# Sex and APOE4-specific links between cardiometabolic risk factors and white matter alterations in individuals with a family history of Alzheimer’s disease

**DOI:** 10.1101/2024.08.21.608995

**Authors:** Stefanie A. Tremblay, R. Nathan Spreng, Alfie Wearn, Zaki Alasmar, Amir Pirhadi, Christine L. Tardif, Mallar M. Chakravarty, Sylvia Villeneuve, Ilana R. Leppert, Felix Carbonell, Yasser Iturria Medina, Christopher J. Steele, Claudine J. Gauthier, PREVENT-AD Research Group

**Affiliations:** Physics department, Concordia University, Montreal, Canada; Montreal Heart Institute, Montreal, Canada; School of Health, Concordia University, Montreal, Canada; Department of Neurology and Neurosurgery, Montreal Neurological Institute, McGill University, Montreal, Canada; Department of Psychiatry, McGill University, Montreal, Canada; McConnell Brain Imaging Centre, Montreal Neurological Institute, Montreal, Canada; Psychology department, Concordia University, Montreal, Canada; Electrical Engineering department, Concordia University, Montreal, Canada; ViTAA Medical Solutions, Montreal, Canada; Department of Biomedical Engineering, McGill University, Montreal, Canada; StoP-AD Centre, Douglas Mental Health Institute Research Centre, Montreal, Canada; Biospective Inc., Montreal, Canada; Ludmer Center for NeuroInformatics and Mental Health, Montreal, Canada; Max Planck Institute for Human Cognitive and Brain Sciences, Leipzig, Germany

**Keywords:** White matter, cardiometabolic risk factors, LDL-cholesterol, sex differences, familial history, APOE4, myelin, memory

## Abstract

**INTRODUCTION:** White matter (WM) alterations are among the earliest changes in Alzheimer’s disease (AD), yet limited work has comprehensively characterized the effects of AD risk factors on WM.

**METHODS:** In older adults with a family history of AD, we investigated the sex-specific and APOE genotype-related relationships between WM microstructure and risk factors. Multiple MRI-derived metrics were integrated using a multivariate approach based on the Mahalanobis distance (D2). The links between WM D2 and cognition were also explored.

**RESULTS:** WM D2 in several regions was associated with high systolic blood pressure, BMI, and glycated hemoglobin, and low cholesterol, in both males and females. APOE4+ displayed a distinct risk pattern, with LDL-cholesterol having a detrimental effect only in carriers, and this pattern was linked to immediate memory performance. Myelination was the main mechanism underlying WM alterations.

**DISCUSSION:** Our findings reveal that combined exposure to multiple cardiometabolic risk factors negatively impacts microstructural health, which may subsequently affect cognition. Notably, APOE4 carriers exhibited a different risk pattern, especially in the role of LDL, suggesting distinct underlying mechanisms in this group.

## 1. BACKGROUND

Recent findings highlight widespread white matter (WM) alterations as a key mechanism in Alzheimer’s disease (AD) development and progression [1–7]. In fact, changes in WM microstructure were found to precede macrostructural atrophy and symptom onset in AD patients [3,4,8], which suggests myelin breakdown is an important contributor to the pathophysiology of AD [9]. The particular vulnerability of oligodendrocytes to various insults (e.g., toxins, oxidative damage) is hypothesized to result in myelin and axonal degeneration over time, precipitating other pathological changes seen in AD such as increased iron [1,9]. Alterations in WM microstructure have thus been suggested as early biomarkers for AD [1,8,10].

Despite WM microstructure being affected early, WM measures are not frequently included in the study of prodromal AD, as more attention has been given to grey matter (GM) abnormalities, such as loss of cortical and hippocampal GM volume [11,12]. Characterizing WM microstructural alterations in individuals at high risk of developing AD is thus crucial in understanding this early stage. Having a familial history and the E4 genotype of the apolipoprotein E (APOE) gene increase the likelihood of developing AD, with a higher risk in females [13]. The APOE4 genotype impacts the brain’s WM microstructure, likely due to its role in the transport of cholesterol, one of the main constituents of myelin [1]. Modifiable risk factors such as physical inactivity, smoking, alcohol consumption, hypertension, diabetes, obesity, and low education also contribute to AD risk [14]. Understanding how these factors impact brain health may inform future interventions.

These modifiable risk factors exhibit complex relationships with WM. For instance, obesity and hypercholesterolemia, known risks for cardiovascular disease and AD [15–17], show mixed associations with WM integrity and cognition [15,16,18]. These inconsistencies may stem from the limited specificity of diffusion MRI measures, typically derived from diffusion tensor imaging (DTI). For instance, reductions in fractional anisotropy (FA), often interpreted as a measure of WM integrity, can be due to axonal loss, but also to increased fiber orientation dispersion [19]. Advanced diffusion models such as NODDI [20,21] and myelin-sensitive techniques such as magnetization transfer imaging are thus needed to fully capture WM microstructural properties [22,23].

The multifaceted interplay between risk factors and WM health may also introduce complexity, leading to seemingly inconsistent results as some factors synergistically influence outcomes while others counteract each other [15,24–27]. Importantly, genetic risk (i.e., APOE4) seems to exacerbate the impact of modifiable risk factors on WM [24–27]. Together, this suggests that the combined effects of multiple risk factors contribute to alterations in WM microstructure. Therefore, integrative approaches, along with advanced WM imaging models, are needed to comprehensively assess the effects of AD risk factors on WM microstructure.

Multi-modal imaging and multivariate frameworks that combine several parameters are promising avenues to harness the complementarity of different neuroimaging-derived metrics [28]. One such approach, the Mahalanobis distance (D2) [29], provides an individual-level measure of deviation relative to a reference group, where voxels with greater D2 values in an individual represent WM areas that differ to a larger extent from the reference group. D2 is a squared distance measure between a point (i.e., measurements in an individual) and a distribution (i.e., reference data) in a multi-dimensional space, integrating several MRI metrics while accounting for covariance between metrics (Fig 1). We previously demonstrated that this method yields an integrative index that meaningfully reflects underlying microstructure in WM in line with known neuroanatomy [30], and that relates to cognitive and motor function in normal subjects [31].

**Figure 1.**
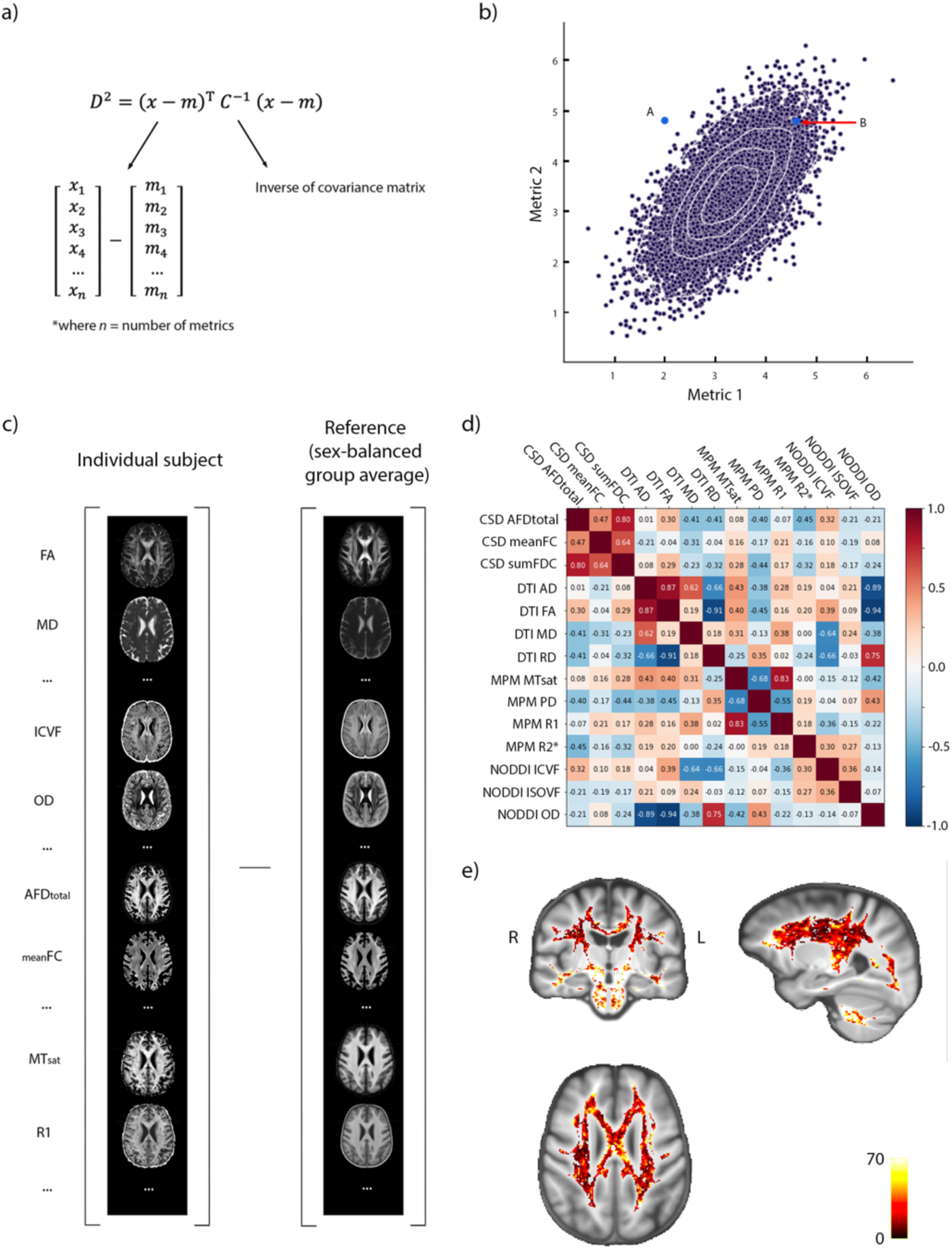
Methodological framework. a) Equation for computing D2. Two vectors, one containing the data of one observation (*x*) and the other containing the mean of all observations for each independent variable (*m*), are subtracted. The covariance between variables is accounted for by multiplying by the inverse of the covariance matrix (*C^-1^*). b) Schematic illustration of the D2 concept in a 2-dimensional space. The purple dots represent the reference distribution (each point represents a subject of the reference group). The probability distance (D2) is the distance, in multivariate space, between each of the blue points (two different subjects; A and B) and the distribution that takes into account the covariance structure of the data. In this example, because the two metrics are positively correlated, A will have a larger D2 value than B. c) The vectors of data are illustrated: the first contains the data of one subject (*x*) and the second contains the reference group average (sex-balanced group) for each metric (*m*). d) The correlation matrix shows relationships between MRI metrics, highlighting the importance of accounting for covariance between variables in multivariate frameworks. e) Example D2 map of a subject. The intensity indicates the amount of deviation in the WM microstructure of this subject compared to the reference, at each voxel.

In this study, we computed voxel-wise deviations in WM microstructure (WM D2) in a cohort of older adults with a family history of AD. We characterized the relationships between known risk factors for AD (education, BMI, blood pressure, cholesterol, and HbA1c) and WM D2 in each sex. The effect of APOE4 genotype on the relationships between risk factors and WM microstructure was also assessed and links with cognition were explored in regions of interest.

## 2. METHODS

### 2.1 Participants

The study population was taken from the PResymptomatic EValuation of Experimental or Novel Treatments for Alzheimer’s Disease (PREVENT-AD) cohort which is composed of older adults (≥ 55 years old) with a familial history of Alzheimer’s disease (parental or multiple-sibling) [32]. The participants, who were followed longitudinally starting in 2011 (some participants are still currently being followed), were all cognitively unimpaired (MoCA ≥ 25, or considered normal after an exhaustive neuropsychological evaluation if < 25, and CDR = 0) at the time of recruitment. Participants gave informed written consent before participating in the study. The procedures of the PREVENT-AD study were approved by the McGill institutional review board and/or Douglas Mental Health University Institute Research Ethics Board. The study was performed in accordance with the ethical standards of the 1964 Declaration of Helsinki.

In this study, we used the ‘stage 2’ MRI data acquired in 2019-2020 (data release 6.0) with a novel imaging protocol that includes multi-shell diffusion-weighted imaging (DWI) and multi-parametric mapping (MPM). Participants who had all DWI and MPM data were included in this cross-sectional study (N= 134). Of those, 97 were female (age = 67.7 ± 4.8 years, education years = 15.3 ± 3.5) and 37 were male (age = 68.6 ± 6.5, education years = 15.7 ± 3.3). Previous time points were not used in this study since these advanced imaging protocols were not acquired in ‘stage 1’ [32].

### 2.2 MRI Protocol

MRI data were acquired on a 3T Siemens PrismaFit scanner at the Douglas Research Centre. The multi-shell DWI sequence was a spin-echo EPI sequence (TR = 3000 ms, TE = 66 ms, phase-encoding direction = posterior-anterior (PA), resolution = 2 mm isotropic) with 100 measurements (isotropically spaced around a sphere) across 3 diffusion-weighted shells with gradient strengths of b = 300 s/mm2 (7 volumes), b = 1000 s/mm2 (29 volumes) and b = 2000 s/mm2 (64 volumes) and 9 volumes acquired without diffusion weighting (b = 0). Five non-diffusion weighted volumes (b = 0) were also acquired in the opposite phase encoding direction (AP) for distortion correction.

An MPM acquisition was performed using three multi-echo gradient echo sequences (resolution = 1 mm isotropic) with different repetition times (TR) and flip angles (α) to obtain images with predominant T1-(TR = 18 ms, 6 echoes, TE = 2.16-14.81 ms, echo-spacing = 2.53 ms, α = 20°), PD-(TR = 27ms, 8 echoes, TE = 2.04-22.20 ms, echo-spacing = 2.57 ms, α = 6°), and MT-weighting (TR = 27 ms, 6 echoes, TE = 2.04-14.89 ms, echo-spacing = 2.57 ms, α = 6°). An off-resonance MT pulse (off-resonance frequency = 2.2 kHz, duration = 12.8 ms, flip angle = 540°) was applied prior to RF excitation to obtain MT-weighting [22]. The RF transmit field was measured using two Siemens turbo-flash sequences with flip angles of 8° and 80° (TR = 5000 ms, TE = 1.83 ms, resolution = 4 x 4 x 16 mm) preceded by a slice-selective preconditioning radiofrequency pulse, yielding anatomical and flip angle maps [35]. RF receive field inhomogeneities were estimated using a pair of PD-weighted turbo-flash sequences acquired using either a body coil or a 32-channel head coil (TR = 344 ms, TE = 1.55 ms, α = 3°, resolution = 2 mm isotropic). The MPM sequence was developed and provided by the McConnell Brain Imaging Centre of The Neuro.

A T1-weighted anatomical scan was also acquired using a Magnetization-Prepared Rapid Acquisition Gradient Echo (MPRAGE) sequence (TR = 2300 ms, TE = 2.96 ms, TI = 900 ms, α = 9°, resolution = 1 mm isotropic) during the same session.

### 2.3 Preprocessing

We computed 14 microstructural metrics from the DWI and MPM data of the ‘stage 2’ time point in 134 participants of the PREVENT-AD cohort. These metrics were derived from the diffusion tensor imaging (DTI) model, the fixel-based analysis framework that derives fibre density and cross-section from fibre orientation distributions (FODs) computed using multi-tissue constrained spherical deconvolution (CSD) [36], and the neurite orientation dispersion and density imaging (NODDI) model [20]. MPM was used to compute quantitative maps of longitudinal relaxation rate (R1), effective transverse relaxation rate (R2*), effective proton density (PD*), and magnetization transfer saturation (MTsat) [37].

#### Diffusion Tensor Imaging

Most processing steps were performed using the MRtrix3 toolbox [38]. DWI data were denoised and then preprocessed using the `dwifslpreproc` Mrtrix3 function, which includes correction for motion and Eddy currents (Eddy tool in FSL 6.0.1), and correction for susceptibility-induced distortions (topup tool in FSL) using b0 volumes of opposite phase-encoding polarities (AP). Preprocessed DWI data were then upsampled to the MPRAGE T1w image resolution (1mm isotropic). Bias field correction was performed using the N4 algorithm of ANTs (3.0) within a mask computed using the brain extraction tool (`bet`) of FSL on the b = 0 preprocessed volume [39]. A brain extraction of all DWI volumes was then applied using the b = 0 mask to remove all non-brain voxels. The tensor was computed on the bias field-corrected DWI data (using `dwi2tensor`) and DTI metrics were then calculated (FA, MD, AD and RD) using `tensor2metric` [40].

#### Fixel-based analysis

The fixel-based analysis (FBA) <u>pipeline</u> which allows the computation of fibre density and cross-section from FODs was followed [38]. First, MPRAGE T1-w images were segmented using the `5ttgen` FSL function of Mrtrix3, which uses the FAST algorithm [41]. Response functions for each tissue type were then computed from the preprocessed DWI data (without bias field correction) and the five-tissue-type (5tt) image using the `dwi2response` function (`msmt_5tt` algorithm) [36]. The bias-uncorrected DWI data was used because bias field correction is performed at a later step in the FBA pipeline [42]. The WM, GM and CSF response functions were then averaged across all participants, resulting in a single response function for each of the three tissue types. Multi-shell multi-tissue CSD was then performed based on the average response functions to obtain an estimation of orientation distribution functions (ODFs) for each tissue type [36]. This step is performed using the `dwi2fod msmt_csd` function of Mrtrix3 within the previously generated brain mask. Bias field correction and global intensity normalization, which normalizes signal amplitudes to make subjects comparable, were then performed on the ODFs, using the `mtnormalise` function in Mrtrix3 [43].

#### Registration

In order to optimize the alignment of WM as well as gray matter, multi-contrast registration was performed. Population templates were generated from the WM, GM and CSF FODs and the brain masks of all participants using the population_template function of Mrtrix3 (with regularization parameters: nl_update_smooth= 1.0 and nl_disp_smooth= 0.75), resulting in a group template for each of the three tissue types [38].

Subject-to-template warps were computed using `mrregister` in Mrtrix3 with the same regularization parameters and warps were then applied to the brain masks, WM FODs, and DTI metrics (i.e., FA, MD, AD and RD) using `mrtransform` [44]. WM FODs were transformed but not reoriented at this step, which aligns the voxels of the images but not the fixels (“fibre bundle elements”). A template mask was computed as the intersection of all warped brain masks (`mrmath min` function). This template mask includes only the voxels that contain data in all subjects. The WM volumes of the five-tissue-type (5tt) 4-D images were also warped to the group template space and averaged across participants to be used as a WM mask for analyses (thresholded at a later step).

#### Computing fixel metrics

The WM FOD template was segmented to generate a fixel mask using the `fod2fixel` function [45,46]. This mask determines the fiber bundle elements (i.e., fixels), within each voxel of the template mask, that will be considered for subsequent analyses. Fixel segmentation was then performed from the WM FODs of each subject using the `fod2fixel` function, which also yields the apparent fibre density (FD) metric. The `fixelreorient` and `fixelcorrespondence` functions were then used to ensure subjects’ fixels map onto the fixel mask [38].

The fibre bundle cross-section (FC) metric was then computed from the warps generated during registration (using the `warp2metric` function) as FC is a measure of how much a fiber bundle has to be expanded/contracted for it to fit the fiber bundles of the fixel template. Lastly, a combined metric, fibre density and cross-section (FDC), representing a fibre bundle’s total capacity to carry information, was computed as the product of FD and FC.

#### Transforming fixel metrics into voxel space

In order to integrate all metrics into the same multi-modal model, fixel metric maps were transformed into voxel-wise maps. As a voxel aggregate of fiber density, we chose to use the *l*=0 term of the WM FOD spherical harmonic expansion (i.e., 1^st^ volume of the WM FOD, which is equal to the sum of FOD lobe integrals) to obtain a measure of the total fibre density (AFDtotal) per voxel. This was shown to result in more reproducible estimates than when deriving this measure from fiber specific FD (i.e., by summing the FD fixel metric) [47]. The FOD *l*=0 term was scaled by the spherical harmonic basis factor (by multiplying the intensity value at each voxel by the square root of 4π).

For the fiber cross-section voxel aggregate measure, we computed the mean of FC, weighed by FD (using the `mean` option of the `fixel2voxel` function). We thus obtained the typical expansion/contraction necessary to align fiber bundles in a voxel to the fixels in the template. Lastly, the voxel-wise sum of FDC, reflecting the total information-carrying capacity at each voxel, was computed using the `fixel2voxel sum` option.

#### NODDI metrics

Bias field corrected DWI data was fitted to the neurite orientation dispersion and density imaging (NODDI) model using the python implementation of Accelerated Microstructure Imaging via Convex Optimization (AMICO) [20,48]. A diffusion gradient scheme file was then created from the bvecs and bvals files. The response functions were computed for all compartments and fitting was then performed on the unbiased DWI volumes, within the brain mask. The resulting parameters were: the intracellular volume fraction (ICVF, also referred to as neurite density), the isotropic volume fraction (ISOVF), and the orientation dispersion index (OD). NODDI metrics were also warped to group space using the transforms generated previously.

#### Multi-parametric mapping

Multi-echo T1-w, PD-w and MT-w images were processed using the hMRI toolbox (v 0.3.0) in Matlab [37,49]. First, all images including field maps were re-oriented using the “AutoReorient” module. This reorientation is based on rigid-body coregistration of a reference image to an MNI template. The first T1-w echo was coregistered to the avg152T1 SPM canonical template and all other images were reoriented. Quantitative R2*, R1, PD and MTsat maps were computed from the reoriented images and field maps using the “Create hMRI maps” module with default parameters. Corrections for RF sensitivity bias, using measured body and head coil sensitivity maps, and for B1 transmit bias field using the TFL B1 mapping method (requires an anatomical image and a flip angle map) were also performed within the “Create hMRI maps” module. MPM maps were warped to the group space.

### 2.4 Computing multivariate distance metric (D2)

The MVComp toolbox was used to compute D2 from the 14 WM features (FA, AD, RD, MD, AFDtotal, meanFC, sumFDC, ICVF, ISOVF, OD, R2*, R1, PD and MTsat) [30]. The first step in computing D2 is to determine the reference from which the multivariate distance will be calculated. Here, because the sample is unbalanced in terms of sex, a sex-balanced reference was built from the 37 male participants and 37 randomly selected females. Demographic characteristics of the reference group, shown in Table 1, were representative of the full sample in terms of age (68.1 ± 5.9), education, cognitive status, and other risk variables. Group averages were then computed from the reference group (N = 74) for each of the 14 metrics using the `compute_average` function of MVComp. The `norm_covar_inv` function was then used to compute the covariance matrix (s) and its pseudoinverse (pinv_s) from the reference. A figure showing the correlations between MRI metrics was generated using the `correlation_fig` function which uses the covariance matrix (s) to calculate correlations (Fig 1). D2 was then computed within MVComp according to this equation:

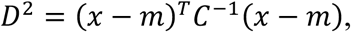

**Table 1.**
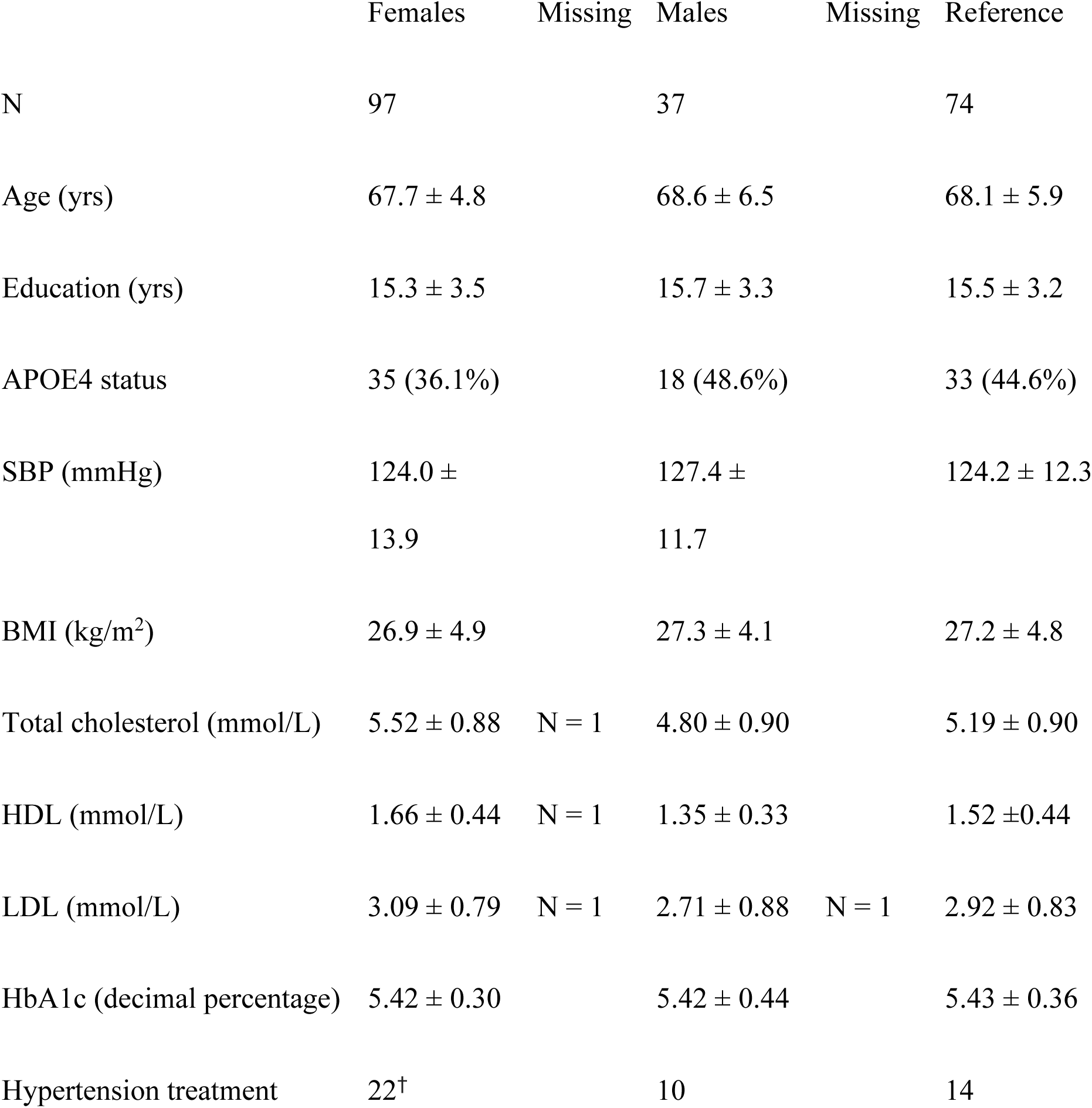
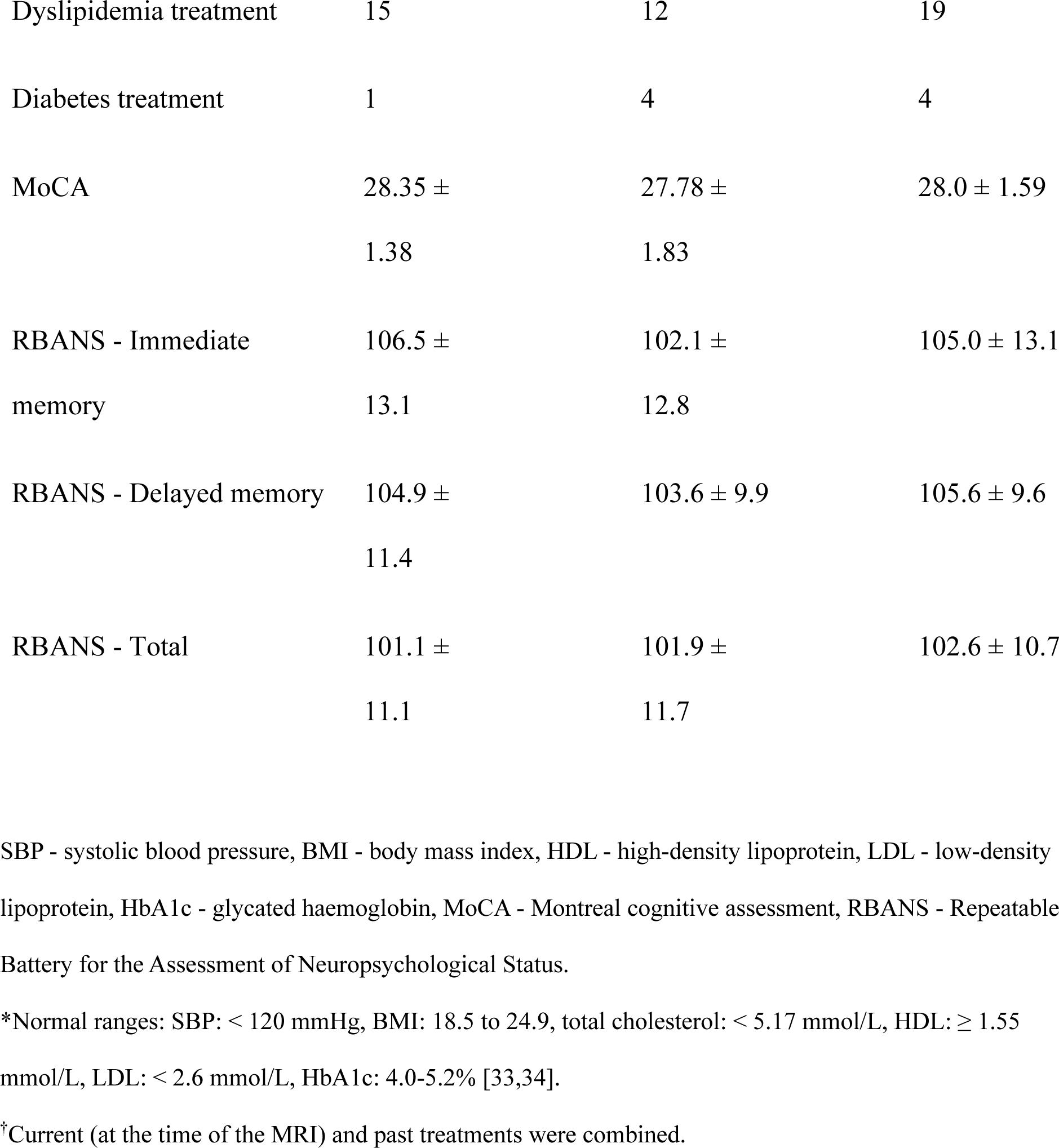
Demographics data for each sex and for the sex-balanced reference group that was used for D2 calculation (mean ± standard deviation). Missing data, if any, is indicated.

where *x* is the vector of data for one observation (e.g., one subject), *m* is the vector of averages of all observations for each independent variable (i.e., MRI metrics), and *C^-1^* is the inverse of the covariance matrix. The `model_comp` function allows the computation of voxel-wise D2 between each subject and the reference average within a specified mask of analysis. Here, a WM mask generated from the average of the WM volumes of the five-tissue-type images of all participants was provided and the threshold was set at 0.99 to limit partial volume effects. The `model_comp` function yields a matrix containing the D2 data of all subjects (of size: number of voxels x number of subjects). The `dist_plot` function was then used to obtain a D2 map (in nifti format) for each subject. The workflow for D2 calculation is illustrated in Figure 1.

The effect of age on D2 was removed by fitting a linear model predicting voxel-wise D2 from age using LinearRegression in sklearn.linear_model and computing the residuals. Residualized D2 data were then normalized using yeo-johnson power transformation voxel-wise in sklearn (version 0.23.2). The residualized and normalized D2 data were used as inputs for the partial least squares (PLS) analysis between WM D2 and risk factors.

Since D2 is a measure of deviation from the reference distribution, the interpretation of D2 depends on the characteristics of the reference sample. High D2 could indicate a region of abnormality if the reference is healthy, or it could be indicative of WM microstructure that is healthier than that of the reference sample if the reference is generally unhealthy. Here, the mean risk variable values of the reference group were slightly higher than the normal healthy ranges for these variables (i.e., SBP, BMI, total cholesterol, LDL, HDL and HbA1c) [33,34], suggesting the latter case.

### 2.5 Blood samples

Blood samples were collected at every annual visit. Variables known to be associated with cardiometabolic risk were used in PLS analyses: total cholesterol, high density lipoprotein (HDL) cholesterol, low density lipoprotein (LDL) cholesterol, and glycated haemoglobin (HbA1c), a clinical index that reflects long-term glycemic control. We used the average of all measurements available in years prior to the MRI date (2011-2018) to reflect cardiometabolic risk history.

### 2.6 Body composition and physiological measures

Blood pressure (BP), heart rate, and body weight (in kg) were measured at every annual visit, while height (in cm) was measured at the eligibility visit. Here we used the average of all measurements available for BP and weight. BMI was calculated as: mean weight (kg) / height^2^ (m). BMI and systolic BP (SBP) were used as ‘risk variables’ in PLS analyses.

### 2.7 APOE4 genotyping

Genotyping methods for this dataset have been described in [32]. Briefly, DNA was isolated from 200 µl whole blood using a QIASymphony apparatus and the DNABlood Mini QIA Kit (Qiagen, Valencia, CA, USA). Allelic variants of AD-related genes including APOE rs429358 and rs7412 were characterized using pyrosequencing (PyroMark24 or PyroMark96) or DNA microarray (Illumina). Participants were classified as either APOE4+ (N= 35 females; N= 17 males) if they had one or more E4 alleles or as APOE4-(N= 61 females; N= 19 males) if they had none. The low number of participants with two E4 alleles (N= 3 females; N= 1 male) did not allow the exploration of a dose-dependent effect of APOE4.

### 2.8 Cognitive assessment

The Repeatable Battery for the Assessment of Neuropsychological Status (RBANS) [50], a brief test (less than 30 minutes) that measures performance in five cognitive domains, was administered to participants at every annual visit. The test yields scaled scores (i.e., age-adjusted index scores with a mean of 100 and standard deviation of 15) for immediate memory, visuospatial/constructional, attention, language and delayed memory. For this study, the RBANS scores of the evaluation conducted on the same day as the MRI session were used and analyses focused on the immediate and delayed memory subscores, the components that have been shown to be the best predictors of AD and mild cognitive impairment [51].

### 2.9 Statistical analyses

#### Relationships between WM microstructure and risk factors

Partial least squares (PLS) analyses were conducted between D2 in WM and risk factors of AD, separately in each sex. PLS is a multivariate statistical approach that can be used to describe spatial relationships between brain MRI data and multiple other variables, in our case risk factors [52]. PLS finds the weight vectors that maximize the covariance between brain data and risk variables, forming new variables called latent variables (LVs). Each WM voxel is assigned a weight, or salience, indicating how strongly it covaries with the pattern of the latent variable, which is a linear combination of the risk factors data.

PLS analyses were conducted in Matlab R2023b (Mathworks Inc.) using the <u>PLS toolbox</u> [52]. The “Regular Behav PLS” was selected as the type of analysis and risk factor data were loaded as the “behavioural data”. Risk factor variables included: education (total number of years of formal education), SBP, BMI, HDL, LDL, total cholesterol, and HbA1c. One participant in each group (males and females) were excluded from these analyses due to missing cholesterol data (see Table 1). The analyses were run with 1000 permutations to determine the significance of each LV, and 1000 bootstraps to determine overall reliability of each voxel’s association to each LV by calculating the standard error of each voxel’s salience value. Only significant LVs (p < 0.05) and voxels with absolute bootstrap ratios (BSR) > 2 (equivalent to p < 0.05) were interpreted. ROIs were created using the `fsl-cluster` function. BSR values greater than the level of significance were used as thresholds for cluster creation to limit the spatial extent of ROIs (described in the Results section).

Similar analyses were conducted between D2 in WM and the same risk factors, this time disaggregating by APOE4 status (APOE4+ if one E4 allele or more; APOE4-if no E4 allele), irrespective of sex. ROIs were created following a similar process as described above.

Analyses were conducted separately in each group to assess patterns specific to each APOE4 group and to each sex. Patterns common to more than one group were then tested for statistical group differences using a 2×2 ANOVA with sex and APOE4 status as fixed factors and with the brain scores (usc) from a PLS analysis in the overall group as the dependent variable. This allowed us to determine whether the pattern was expressed more strongly in one group compared to the others and to test for interaction between APOE4 and sex.

#### Relationships between deviations in WM microstructure and cognition

Correlation analyses were conducted between mean D2 in ROIs (significant clusters from PLS analyses) and the immediate and delayed memory subscores of the RBANS. Analyses were targeted to these two subscores, known to be the most affected cognitive domains in AD [51], and to significant clusters identified in the PLS analysis, to limit the number of comparisons. Correction for multiple comparisons was performed using the false discovery rate (FDR) Benjamini-Hochberg method (FDR-corrected p-value < 0.05 was considered statistically significant).

#### Determining feature importance in regions of interest

The relative contributions of each feature (i.e., MRI metric) to D2 in significant ROIs (of size > 100 voxels) were then extracted using the `return_raw` option of the `model_comp` function in MVComp. The `return_raw` option yields a matrix of size (number of voxels) x (number of metrics) x (number of subjects). Contributions were then summarised by averaging distance values across voxels within the ROI and across subjects and dividing by the total distance (for all features), resulting in one distance value per metric, expressed as a percentage, for each ROI. This analysis provides a measure of the importance of each metric in determining D2 in the ROI.

## 3. RESULTS

### 3.1 Relationships between WM microstructure and risk factors in each sex

Significant patterns of covariance were found between risk factors for AD and WM D2 in both males and females. In males, only the first LV of the PLS analysis was significant (p = 0.002) and it explained 34.4% of total crossblock covariance (Fig 2a-b). Low SBP, low BMI, low HbA1c and high cholesterol (total chol, HDL and LDL) were associated with high D2 in several WM regions including the body of corpus callosum, superior corona radiata, superior thalamic radiation (bilaterally) and the right frontal aslant tract. In females, only the first LV was significant (p < 0.001) and explained 40.9% of total crossblock covariance (Fig 2c-d). Similar relationships were observed but in slightly different WM regions. D2 in the superior longitudinal fasciculus, corticospinal tract, cingulum, splenium of corpus callosum, posterior corona radiata, and arcuate fasciculus (bilaterally), as well as the right forceps major and body of corpus callosum was associated with these risk factors in females. Generally, associations were found in more frontal and parietal regions in males, while they were found in more posterior and temporal locations in females. There were also overlapping regions in both sexes, specifically in parietal regions and in WM tracts underlying the precentral gyrus. Education was the only non-significant factor in both groups. Figure 2 shows the strength and direction of the relationships between D2 and each risk factor (left panel), as well as the WM regions in which those relationships are located (right panel). Only significant voxels (|BSR| > 2.0) and those belonging to clusters of size > 100 voxels are shown. Clusters were formed from significant voxels. Thresholds higher than the significance limit (|BSR| = 2.0, equivalent to p = 0.05) were used to limit the spatial extent of clusters and different cluster thresholds were used in each sex to result in similarly sized clusters, so that D2 was averaged across a similar number of voxels for analyses with cognition (|BSR| > 2.5 in females and > 3.0 in males). Further analyses were focused on clusters of size > 100 voxels. Cluster information, including the size, maximal |BSR| and brain region, is displayed in Table 2.

**Figure 2.**
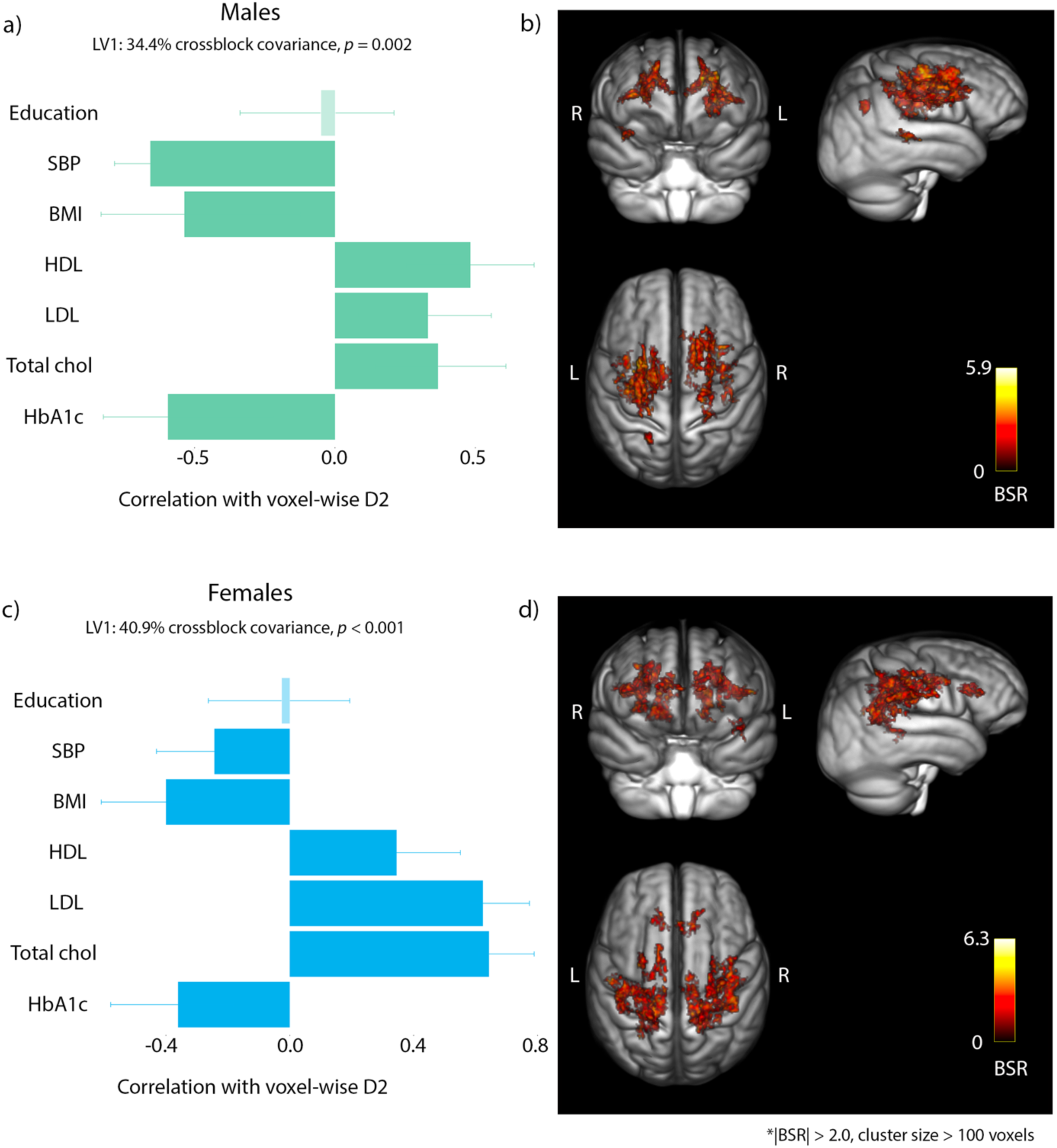
Relationships between D2 in WM and risk factors in each sex. Left panel (a & c): The strength and direction of the relationship that each risk factor has with D2 in the voxels shown on the brain images on the right. Error bars show 95% confidence intervals. Correlations are non-significant when confidence intervals overlap with zero (faded bar). Right panel (b & d): Colored voxels (|BSR| > 2.0) have a positive relationship with the patterns shown in the left panel. The BSR maps are overlaid on a MPRAGE T1w group average image. Males (a-b) Several risk factors were associated with D2 across broad WM regions. Higher D2 was associated with lower SBP, BMI and HbA1c and with higher HDL, LDL and total cholesterol. Females (c-d) Similar relationships were observed in females but across different WM regions.

**Table 2.**
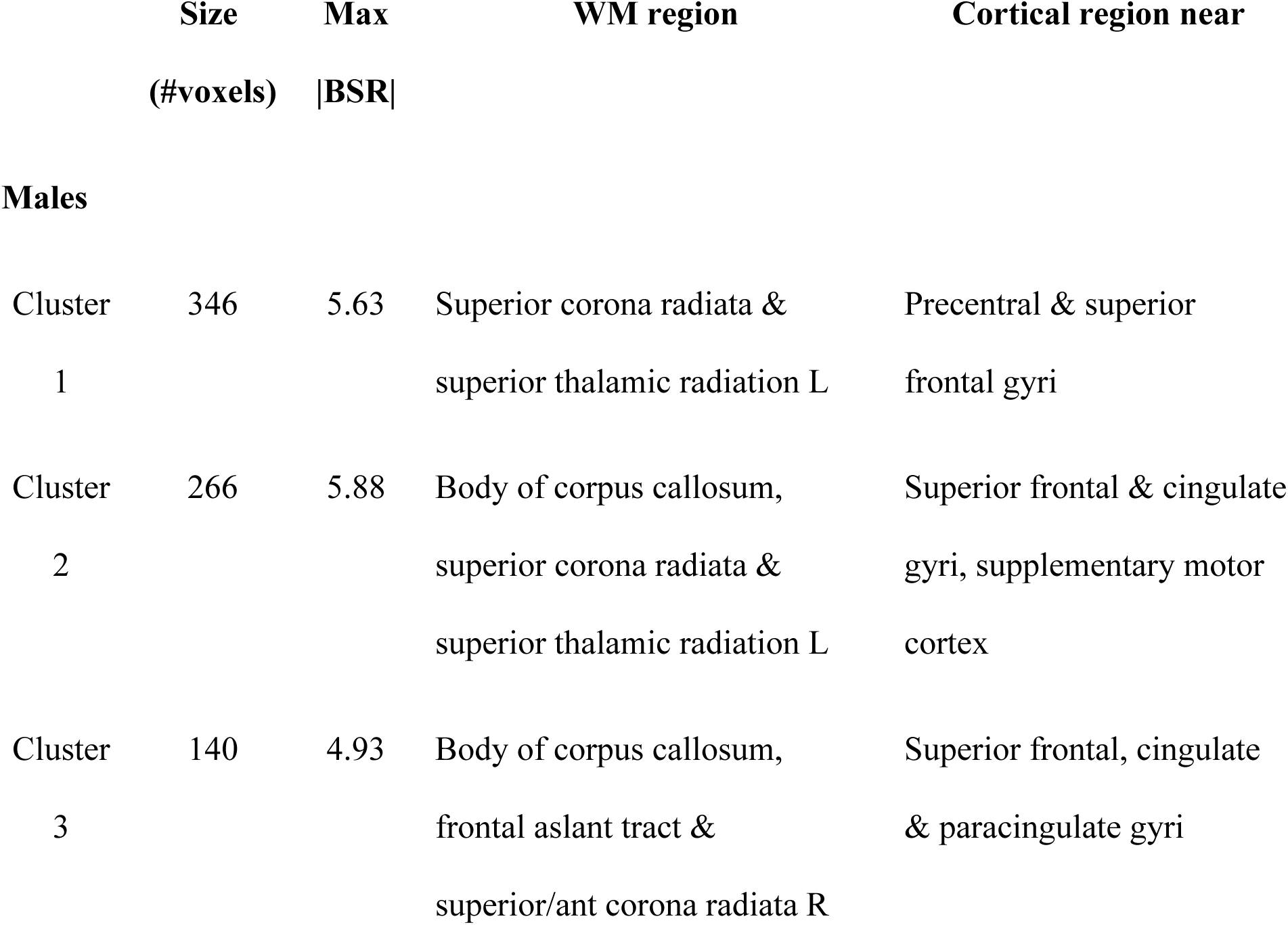
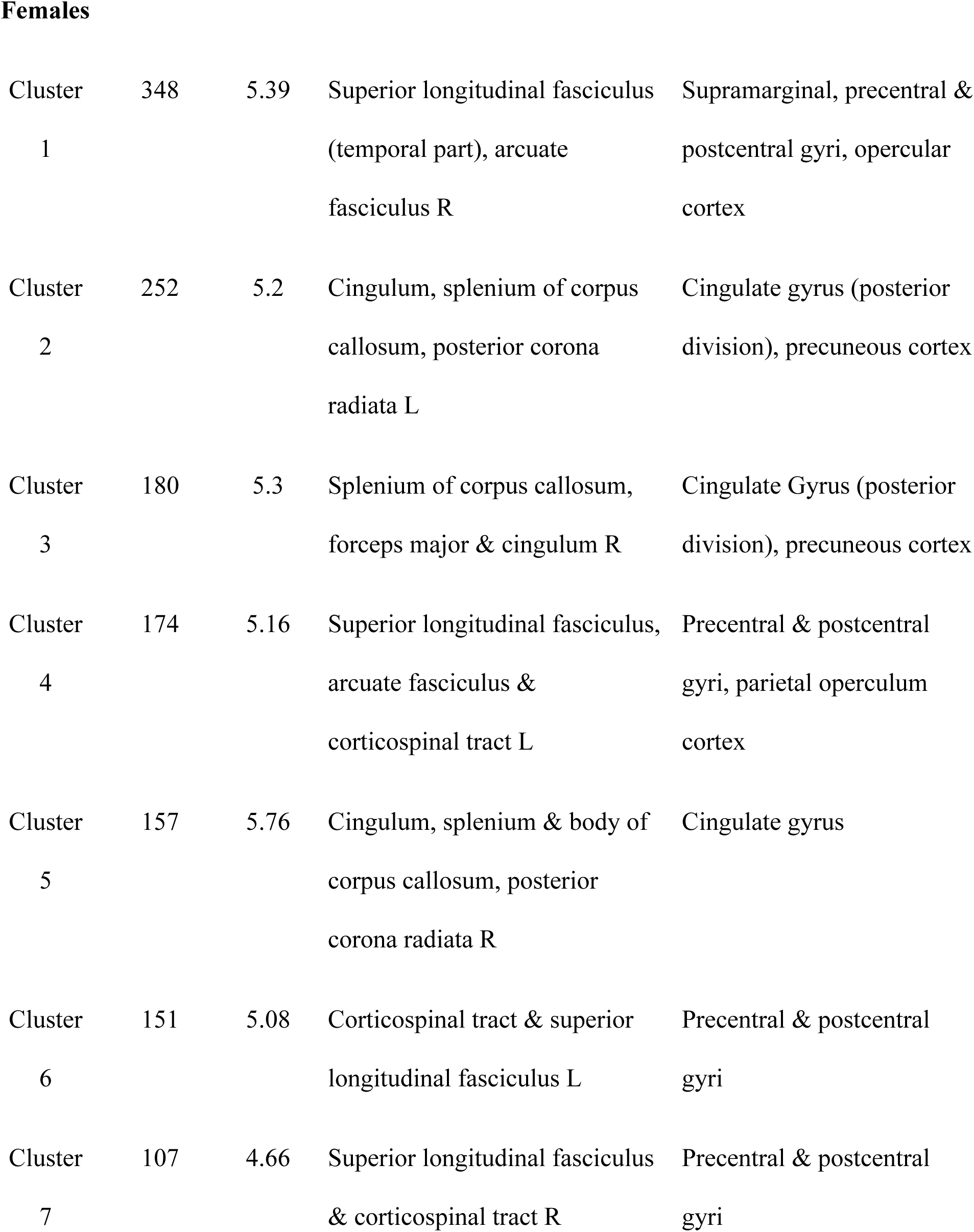
Cluster information (PLS analyses in each sex). Location identified according to the JHU ICBM-DTI-81 White-Matter and XTRACT HCP Probabilistic tract atlases and cortical region closest to the WM region identified using the Harvard-Oxford cortical structural atlas.

|  | Size<br>(#voxels) | Max<br>\|BSR\| | WM region | Cortical region near |
| --- | --- | --- | --- | --- |
| **Males** |  |  |  |  |
| Cluster 1 | 346 | 5.63 | Superior corona radiata & superior thalamic radiation L | Precentral & superior frontal gyri |
| Cluster 2 | 266 | 5.88 | Body of corpus callosum, superior corona radiata & superior thalamic radiation L | Superior frontal & cingulate gyri, supplementary motor cortex |
| Cluster 3 | 140 | 4.93 | Body of corpus callosum, frontal aslant tract & superior/ant corona radiata R | Superior frontal, cingulate & paracingulate gyri |
| Females |  |  |  |  |
| Cluster 1 | 348 | 5.39 | Superior longitudinal fasciculus (temporal part), arcuate fasciculus R | Supramarginal, precentral & postcentral gyri, opercular cortex |
| Cluster 2 | 252 | 5.2 | Cingulum, splenium of corpus callosum, posterior corona radiata L | Cingulate gyrus (posterior division), precuneous cortex |
| Cluster 3 | 180 | 5.3 | Splenium of corpus callosum, forceps major & cingulum R | Cingulate Gyrus (posterior division), precuneous cortex |
| Cluster 4 | 174 | 5.16 | Superior longitudinal fasciculus, arcuate fasciculus & corticospinal tract L | Precentral & postcentral gyri, parietal operculum cortex |
| Cluster 5 | 157 | 5.76 | Cingulum, splenium & body of corpus callosum, posterior corona radiata R | Cingulate gyrus |
| Cluster 6 | 151 | 5.08 | Corticospinal tract & superior longitudinal fasciculus L | Precentral & postcentral gyri |
| Cluster 7 | 107 | 4.66 | Superior longitudinal fasciculus & corticospinal tract R | Precentral & postcentral gyri |

### 3.2 Relationships between WM microstructure and risk factors in each APOE4 group

Different patterns of covariance were found between risk factors for AD and WM D2 in the APOE4+ and APOE4-groups. In APOE4+, LV1 (p = 0.013, crossblock covariance = 34.9%) and LV2 (p = 0.048, crossblock covariance = 22.5%) were significant. The LV1 pattern revealed that low BMI and high cholesterol (total chol, HDL and LDL) were associated with high D2 in several WM regions including the left body of corpus callosum, frontal aslant tract, superior thalamic radiation, and arcuate fasciculus (Fig 3a-b). These risk factors and their directions of association to D2 represent a subset of the pattern seen in sex-specific analyses. On the other hand, LV2 revealed a different risk pattern: low SBP, low BMI, high HDL, low LDL, and low HbA1c were associated with high D2 in the right superior longitudinal fasciculus, superior corona radiata, superior thalamic radiation, and corticospinal tract (Fig 3c-d). Generally, associations of the first LV were found in the left hemisphere and included commissural fibers such as the corpus callosum, while LV2 associations were found mostly on the right and included projection fibers such as the superior corona radiata as well as association tracts.

**Figure 3.**
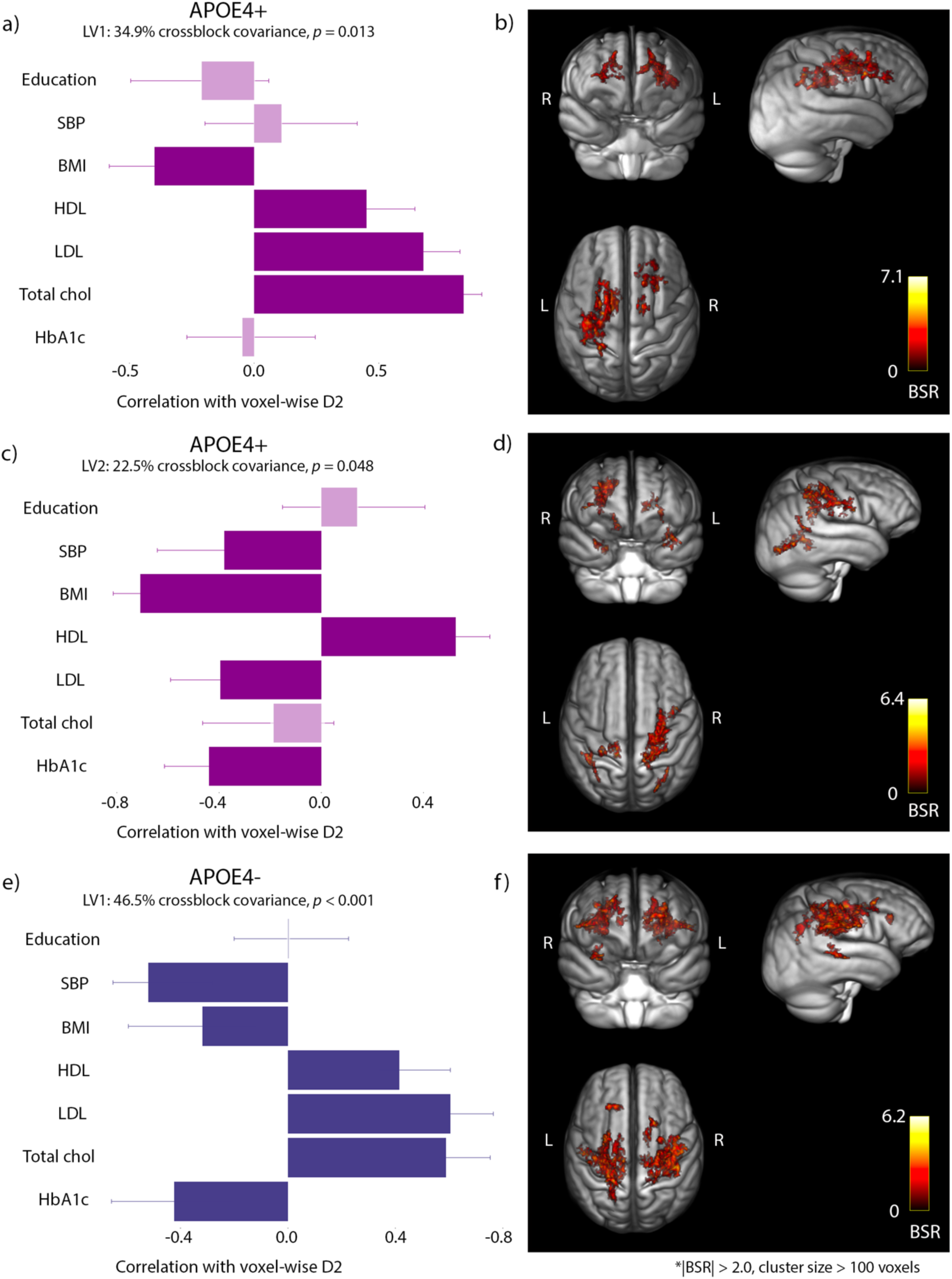
Relationships between D2 in WM and risk factors in each APOE4 group. Left panel: The strength and direction of the relationship that each risk factor has with D2 in the voxels shown on the brain images on the right. Error bars show 95% confidence intervals. Correlations are non-significant when confidence intervals overlap with zero (faded bar). Right panel: Colored voxels (|BSR| > 2.0) have a positive relationship with the patterns shown in the left panel. The BSR maps are overlaid on a MPRAGE T1w group average image. **APOE4+** (a-b) LV1: Higher D2 was associated with lower BMI and higher HDL, LDL and total cholesterol. (c-d) LV2: Higher D2 was associated with low SBP, low BMI, high HDL, low LDL, and low HbA1c. **APOE4-** (e-f) Higher D2 was associated with low SBP, BMI, HbA1c and with high HDL, LDL and total cholesterol.

In APOE4-, only the first LV of the PLS analysis was significant (p < 0.001) and it explained 46.5% of total crossblock covariance (Fig 3e-f). The risk factors pattern was very similar to that of previous analyses (sex-disaggregated PLS analyses). Low SBP, low BMI, low HbA1c and high cholesterol (total chol, HDL and LDL) were associated with high D2 in broad WM regions including the superior longitudinal fasciculus, arcuate fasciculus, superior corona radiata, and corticospinal tract. Significant regions of the PLS analysis in APOE4-overlapped to a large extent with significant regions seen in sex analyses.

Education was non-significant in all LVs. Figure 3 shows the strength and direction of the relationships between D2 and each risk factor (left panel), as well as the WM regions in which those relationships are located (right panel). Only significant voxels (|BSR| > 2.0) and those belonging to clusters of size > 100 voxels are shown. Clusters were formed from significant voxels. Thresholds of |BSR| > 2.5 in APOE4+ and > 3.0 in APOE4-were used for clustering to result in similar size clusters across groups and further analyses were focused on clusters of size > 100 voxels. Cluster information is displayed in Table 3.

**Table 3.**
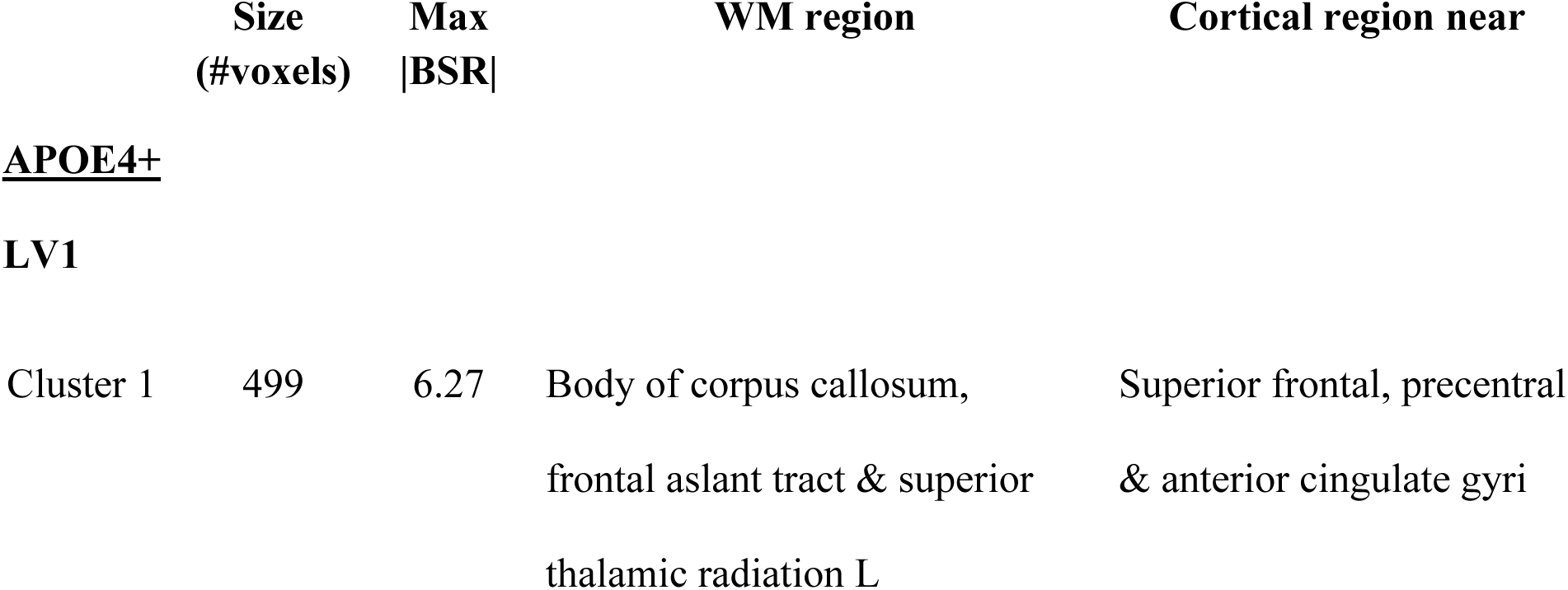
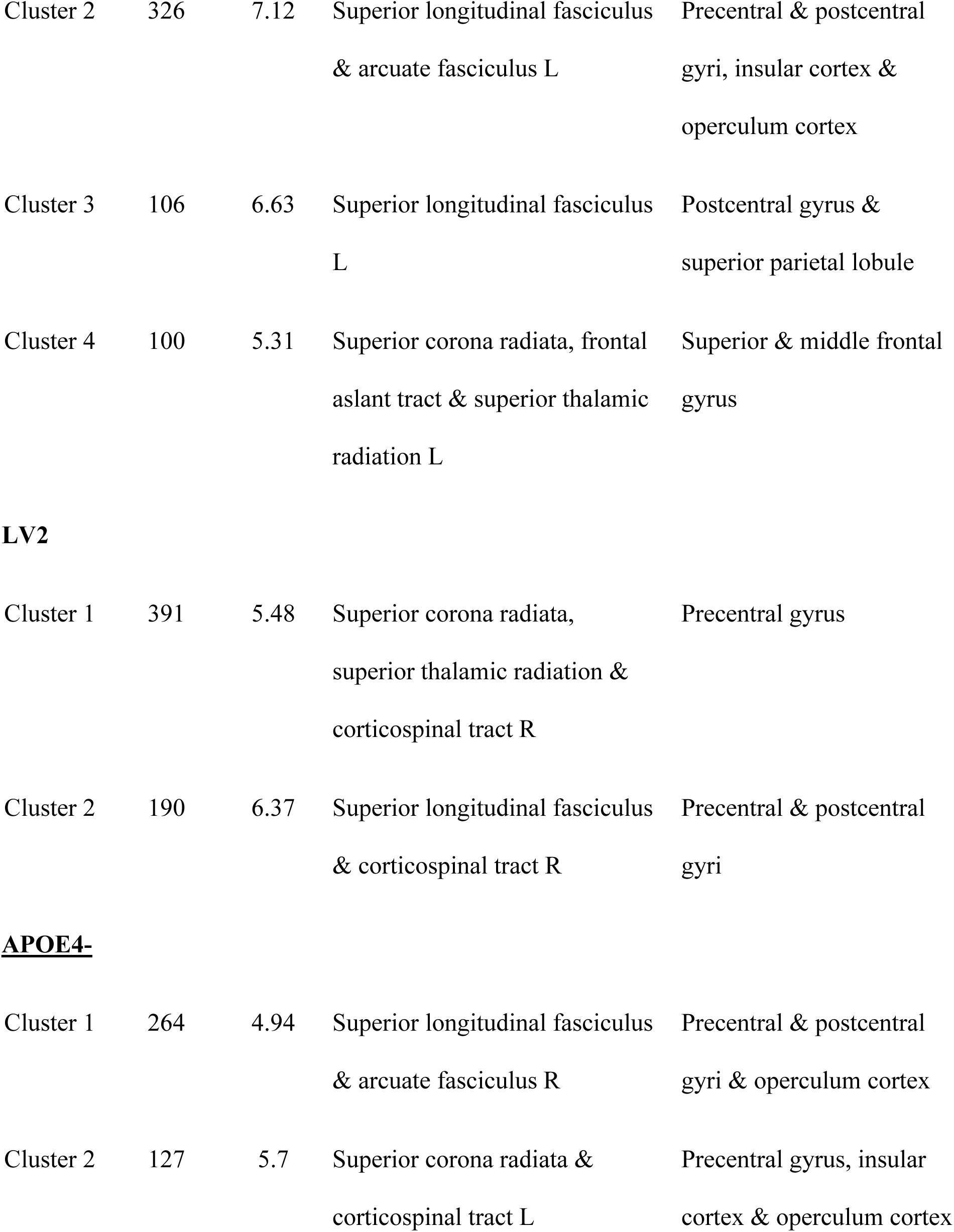
Cluster information (PLS analyses in each APOE4 group). Location identified according to the JHU ICBM-DTI-81 White-Matter and XTRACT HCP Probabilistic tract atlases and cortical region closest to the WM region identified using the Harvard-Oxford cortical structural atlas.

|  | Size<br>(#voxels) | Max<br>\|BSR\| | WM region | Cortical region near |
| --- | --- | --- | --- | --- |
| **<u>APOE4+</u>** |  |  |  |  |
| **LV1** |  |  |  |  |
| Cluster 1 | 499 | 6.27 | Body of corpus callosum,<br>frontal aslant tract & superior<br>thalamic radiation L | Superior frontal, precentral<br>& anterior cingulate gyri |
| Cluster 2 | 326 | 7.12 | Superior longitudinal fasciculus & arcuate fasciculus L | Precentral & postcentral gyri, insular cortex & operculum cortex |
| Cluster 3 | 106 | 6.63 | Superior longitudinal fasciculus L | Postcentral gyrus & superior parietal lobule |
| Cluster 4 | 100 | 5.31 | Superior corona radiata, frontal aslant tract & superior thalamic radiation L | Superior & middle frontal gyrus |
| **LV2** |  |  |  |  |
| Cluster 1 | 391 | 5.48 | Superior corona radiata, superior thalamic radiation & corticospinal tract R | Precentral gyrus |
| Cluster 2 | 190 | 6.37 | Superior longitudinal fasciculus & corticospinal tract R | Precentral & postcentral gyri |
| **APOE4-** |  |  |  |  |
| Cluster 1 | 264 | 4.94 | Superior longitudinal fasciculus & arcuate fasciculus R | Precentral & postcentral gyri & operculum cortex |
| Cluster 2 | 127 | 5.7 | Superior corona radiata & corticospinal tract L | Precentral gyrus, insular cortex & operculum cortex |

Because a general common pattern was observed in both sexes and in the APOE4-group, we performed another PLS analysis in the whole sample to test for group differences and interactions between sex and APOE4. This analysis showed a very similar pattern as that observed in these groups (Supplementary Fig 1). The ANOVA on the brain scores (i.e., usc) from this analysis revealed a significant main effect of sex (p < 0.001), indicating that males expressed the pattern of the LV more strongly than females (Supplementary Table 1 and Supplementary Fig 2). There were no significant APOE4 group differences and no significant sex x APOE4 interaction (p > 0.05) (Supplementary Table 1).

### 3.3 Determining feature importance in regions of interest

In females, features’ contributions were extracted in the 7 clusters that were significant in the PLS analysis (Fig 4a). D2 in cluster 1 was driven mainly by MTsat (28.8%), R1 (23.4%), and RD (12.5%). In cluster 2, meanFC (29.9%), R1 (23.6%), and MTsat (11.9%) contributed the most to D2. Similarly, in cluster 3, meanFC (28.1%), R1 (18.3%), and MTsat (14.3%) were the top contributors. In cluster 4, R1 (28.7%), MTsat (25.5%), and ISOVF (15.6%) contributed most to D2. D2 in cluster 5 was driven mainly by R1 (44.5%) and meanFC (23.2%). In cluster 6, R1 (20.8%), ISOVF (18.1%), meanFC (16.0%), and MTsat (14.2%) were the metrics that contributed the most to D2. In cluster 7, D2 was mainly driven by R1 (25.3%), meanFC (19.5%), AD (13.5%), and MD (10.8%). Overall, R1, meanFC, and MTsat were the most important metrics in females. The isotropic volume fraction (ISOVF) metric from NODDI was also an important contributor in 2 clusters.

**Figure 4.**
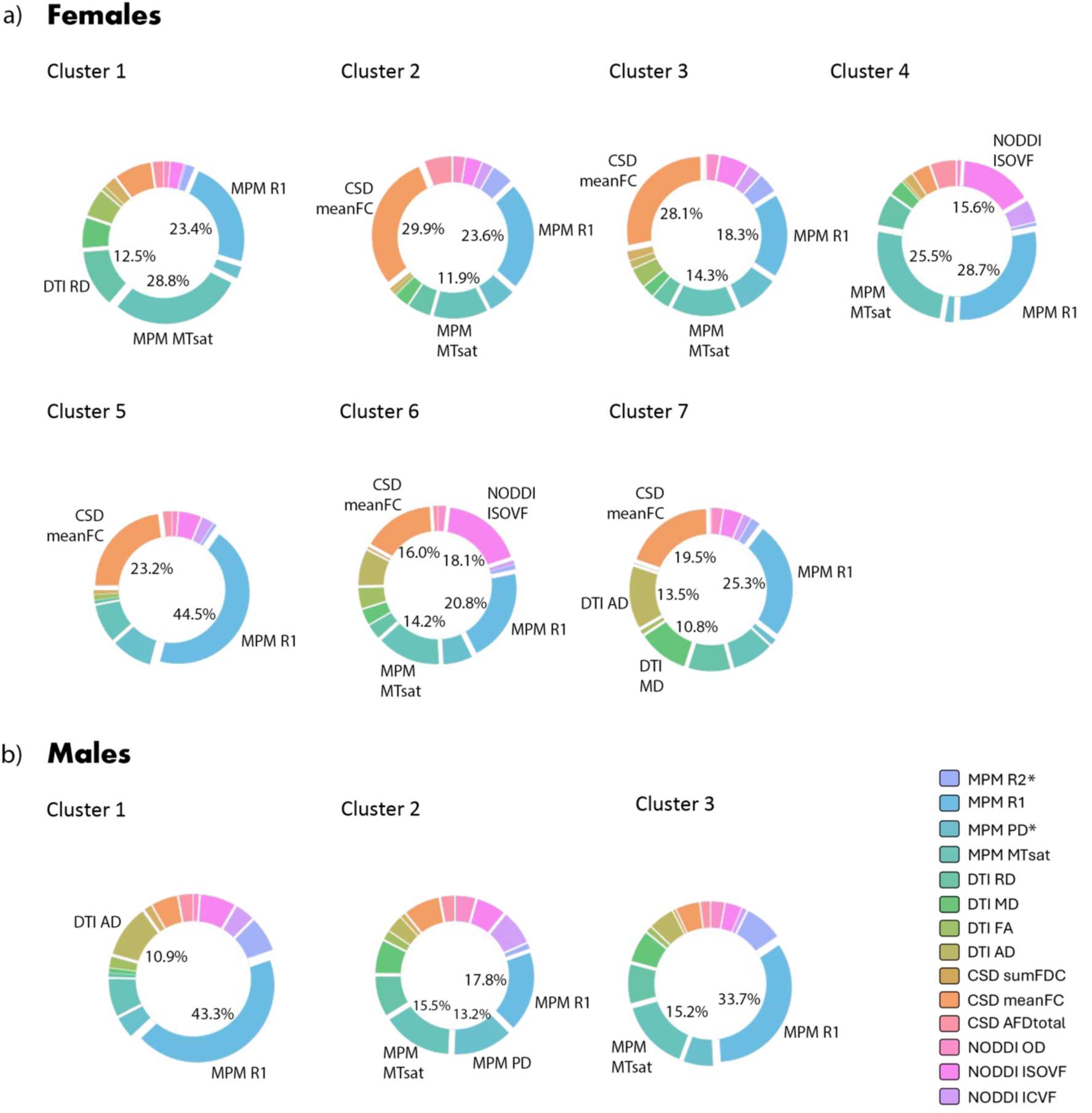
Features contribution to D2 in each significant cluster from PLS analyses in females (a) and males (b). For each significant cluster, the relative contribution (%) of each MRI metric is indicated by its size on the pie chart (see the legend for color of each MRI metric). The metric name and its contribution (in %) is indicated only for the most important contributors (those that account for >10%), for clarity. MPM R1 = macromolecular content (axons and myelin) [53]; MPM MTsat = more specific to myelin content [22]; MPM PD = amount of water (if increased could reflect neurite atrophy); CSD meanFC = fiber bundle cross-section [42]; NODDI ISOVF = amount of free water (if increased it could reflect neurite atrophy) [20]; AD = axonal integrity; RD = myelin integrity [54]; MD = overall diffusivity (typically increased with higher water content/cell atrophy).

In males, features’ contributions were extracted in the 3 significant clusters from PLS analysis (Fig 4b). D2 in cluster 1 was driven mainly by R1 (43.3%) and, to a lesser extent, by AD (10.9%). In cluster 2, R1 (17.8%), MTsat (15.5%), and PD (13.2%) contributed the most to D2. In cluster 3, R1 (33.7%) and MTsat (15.2%) were the top contributors. Like in females, R1 and MTsat emerged as top contributors to D2 in males. However, meanFC was not an important contributor to D2 (<10%) in any of the males’ clusters.

In the APOE4+ group, there were 4 clusters from the first LV and 2 clusters from the second LV (Fig 5a-b). D2 in the first LV1 cluster was driven mainly by R1 (28.8%), MTsat (20.4%), and PD (12.6%). In cluster 2, R1 (28.4%), MTsat (28.1%), and meanFC (10.5%) were the top contributors. In cluster 3, R1, (26.1%), MTsat (17.1%), meanFC (12.5%), and ISOVF (12.4%) contributed most to D2. D2 in cluster 4 was driven mainly by R1 (36.6%) and meanFC (20.5%). In cluster 1 of the second LV, R1 (26.0%), MTsat (23.3%), and meanFC (18.5%) contributed most to D2. In cluster 2, MTsat (25.7%), R1 (24.4%), and OD (11.2%) were the metrics that contributed most to D2.

**Figure 5.**
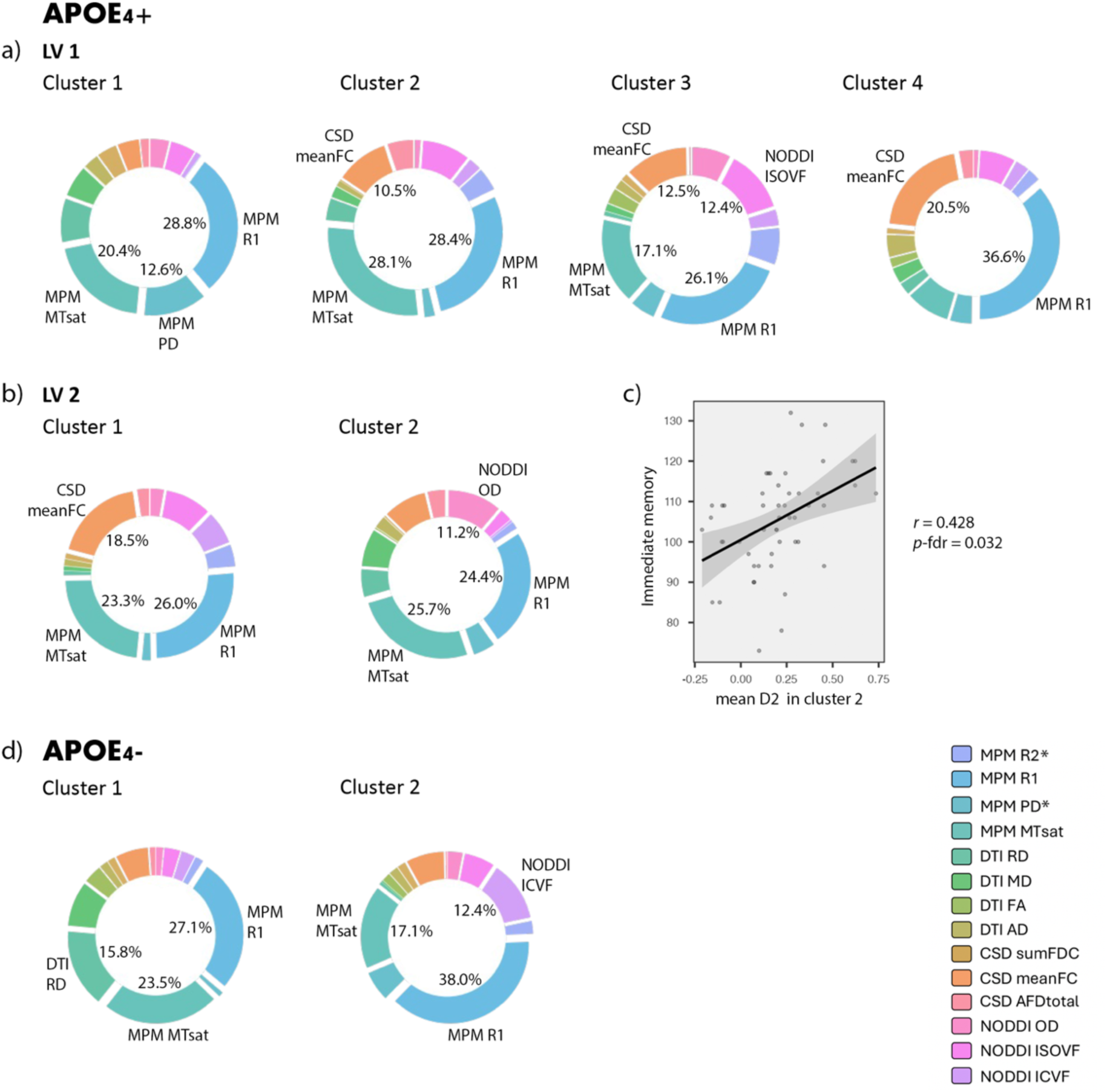
Features contribution to D2 in each significant cluster from PLS analyses in APOE4+ (a-b) and APOE4-(d). For each significant cluster, the relative contribution (%) of each MRI metric is indicated by its size on the pie chart (see the legend for color of each MRI metric). The metric name and its contribution (in %) is indicated only for the most important contributors (those that account for >10%), for clarity. c) Plots are shown for significant correlations between D2 and the RBANS memory items. Immediate memory was positively associated with D2 in cluster 2 of the APOE4+ analysis (LV2). MPM R1 = macromolecular content (axons and myelin) [53]; MPM MTsat = more specific to myelin content [22]; MPM PD = amount of water (if increased could reflect neurite atrophy); CSD meanFC = fiber bundle cross-section [42]; NODDI ISOVF = amount of free water (if increased it could reflect neurite atrophy); NODDI ICVF = neurite density; NODDI OD = orientation dispersion of fiber tracts [20]; RD = myelin integrity [54].

In the APOE4-group, features’ importance was extracted in the 2 significant clusters (Fig 5d). D2 in cluster 1 was driven mainly by R1 (27.1%), MTsat (23.5%), and RD (15.8%). In cluster 2, R1 (38.0%), MTsat (17.1%), and ICVF (12.4%) contributed the most to D2.

### 3.4 Relationships between deviations in WM microstructure and cognition

To understand the associations between deviations in WM microstructure and cognition, correlation analyses were performed between D2 in significant clusters from the PLS analysis disaggregated by APOE4 status (8 clusters; Table 3) and scores in the immediate and delayed memory RBANS items. D2 in the two LV2 clusters was positively associated with immediate memory. The first cluster, located in a WM region corresponding to part of the right superior corona radiata, superior thalamic radiation and corticospinal tract, had a product-moment correlation (*Pearson’s*) *r* = 0.313 and a *p* value = 0.024 (*p*-fdr corrected = 0.192). Cluster 2, located in a WM region corresponding to part of the right superior longitudinal fasciculus and corticospinal tract, had a product-moment correlation *r* = 0.428, *p* = 0.002 (*p*-fdr corrected = 0.032) (Fig 5c). Only the correlation in cluster 2 remained significant after FDR correction. All other correlations were non-significant (p > 0.05).

## 4. DISCUSSION

In this study, we identified the presence of WM microstructural impairments linked to cardiometabolic risk factors in individuals with a family history of Alzheimer’s disease (AD). These impairments were identified using a novel approach, whereby we investigated the sex-specific and APOE genotype-related relationships between WM microstructural deviations, quantified using a multivariate score derived from several MRI-derived features, and cardiometabolic risk factors.

### Sex-related effects

In our sex-disaggregated analysis, we found that in both males and females, high systolic blood pressure, high BMI, high HbA1c (blood sugar levels), and low cholesterol (total, HDL, and LDL) were associated with low D2 (Fig 2, left panel). Due to the directions of relationships between risk factors and D2, we inferred that, in our analyses, greater D2 is likely to represent a healthier state. Since D2 represents the amount of deviation from the reference distribution (i.e., sex-balanced reference group), this would mean that higher D2 indicates WM microstructure that is healthier than the average of our reference group. Although the patterns of association were similar in males and females, the WM regions in which these relationships were observed differed between sexes (Fig 2, right panel), with partially overlapping significant clusters, but more frontal regions in males and more posterior and temporal locations in females. There was also a significant sex difference, where males expressed this general risk–WM pattern more strongly than females. There are known sex differences in WM microstructure [55,56] and in the aging trajectory of myelin [57]. Age-related declines in WM health follow an anterior to posterior gradient and these changes start occurring later in life in females, likely owing to the pro-myelinating effects of female hormones [1,57]. In this study, we regressed out age to remove any age effects and the pattern of WM differences we observe does not fit this model. The sex differences observed may thus be due to factors more specifically related to sex differences in WM susceptibility to vascular risk factors. More research is needed to understand the sex-specific relationships between risk factors, early pathological changes and AD, as well as to disentangle the contributions of gender [58], which may confound these relationships.

The directions of several of the associations we found between WM D2 and risk factors are in line with the literature. Several studies report WM alterations in hypertensive individuals and the effects of high blood pressure may start accumulating as early as the fourth decade of life [59]. Obesity has also been associated with changes in WM microstructure such as decreased FA and myelin content (R1) in several WM tracts [60,61]. In line with the extant literature on subclinical hyperglycemia [62,63], we also identified WM differences associated with HbA1c levels, even though the vast majority of participants did not have diabetes (99% of females and 92% of males). Moreover, HDL cholesterol was positively associated with WM D2, in line with the well-established protective role of HDL on cognition and brain structure [64].

### APOE4-dependent effects of LDL

We found complex relationships between WM microstructure and LDL. Peripheral LDL and total cholesterol were positively associated with better WM health (high D2) in all but one latent variable. While LDL and total cholesterol are typically thought of as being detrimental, evidence on their impact on the brain’s WM and on cognition is unclear [15,16,18,65–67]. These discrepancies may be due to the unknown contribution of oxidized LDL to total LDL. Oxidation of LDL, which is enhanced in inflammatory states when oxidative stress is high, has been shown to be a better predictor of atherosclerosis and cardiovascular disease than LDL itself [68–71] and is also associated with deleterious effects on brain health [72,73]. It is thus likely that the relationship we observed between LDL and WM was due to predominantly non-oxidized LDL. The fact that LDL and HDL cholesterol were related to WM D2 in the same direction also supports this hypothesis as the antioxidant property of HDL would contribute to preventing LDL oxidation [64,69,74]. Furthermore, as ∼80% of participants in this study were taking lipid-lowering medications, LDL oxidation may be reduced in the participants taking statins [75].

In contrast, we found that high LDL was associated with poorer WM health in APOE4 carriers (LV2). This is consistent with a study reporting a detrimental effect of elevated LDL on WM microstructure in APOE4 carriers, but a beneficial effect in non-carriers [76]. As a cholesterol-transporter, APOE4 may modulate the impact of LDL on WM microstructure through increased LDL circulation time, increased free radical formation and decreased plasma antioxidant concentrations, increasing LDL oxidation [72]. Overall, our study supports the idea of differential effects of LDL-cholesterol on the brain’s WM depending on APOE genotype. However, future studies that include measurements of oxidized LDL and a larger sample size (especially of APOE4 homozygotes) are needed to confirm these findings.

The distinct pattern observed in APOE4 carriers (LV2), where low D2 was associated with high LDL, low HDL-cholesterol, high HbA1c, high BMI and high SBP, was found to be linked with cognition. D2 in a cluster of this LV was positively associated with immediate memory performance, indicating that this pattern of risk factors likely had a negative impact on cognition in APOE4 carriers. The direction of the relationship with cognition also supports our interpretation of low D2 reflecting poor WM health. On the other hand, in non-carriers, WM D2 in regions associated with risk factors did not relate with cognition. Together, this suggests that WM health is differentially affected by cardiometabolic risk factors in APOE4 carriers and that the pattern uncovered by LV2 may be more detrimental to cognitive health.

### Role of myelin and other components

Several WM regions were associated with the patterns of risk factors discussed above and extracting the contribution of each MRI feature to D2 in these regions revealed that inter-individual variations in myelin content (as measured by R1 and MTsat) was a major contributor in most significant clusters. Our results are partially in line with the myelin breakdown theory stating that late-myelinating WM tracts would be especially vulnerable to aging and adverse risk factors such as those investigated in this study [1,2,9,77]. For instance, decreased FA in late-myelinating tracts has been reported in individuals with elevated glycated hemoglobin (HbA1c) [25]. Most of our significant clusters were located in late-myelinating regions (i.e., supramarginal, superior frontal, superior parietal, superior temporal, and precuneus WM), but we also found significant associations in the splenium of the corpus callosum, a region that develops at an intermediate stage [1,2,9,25,77].

### Strengths and limitations

In this study, we assessed relationships between risk factors and WM microstructure separately in each sex and APOE4 group, which allowed the identification of patterns specific to APOE4 carriers. Importantly, this pattern would not have been detected in a whole sample analysis (see Supplementary Fig 1 and Table 1). Another strength of our study is the use of a multivariate approach to integrate several MRI measures of WM, allowing for a comprehensive assessment of the biological mechanisms underlying WM differences [30]. This is of interest because multiple pathological mechanisms (e.g., demyelination, axonal changes, iron accumulation) are likely involved concurrently in AD and in its prodromal stage [78]. However, the D2 method has some inherent limitations. Because D2 is a squared measure, the directionality of differences is non-specific [30]. Future studies could address this limitation by integrating models of ground-truth biophysical properties to better interpret these differences, or by stratifying groups based on the expected direction of change to have a strong prior on the directions of deviations. Further, the high sensitivity of D2 makes it susceptible to registration inaccuracies and partial voluming. Special attention must thus be paid to optimize alignment and minimize partial voluming. In this study, strict masking (i.e., 0.99 of group average tissue segmentation) was applied to restrict the analyses to voxels containing only WM. Another limitation of this study is that the sample size did not allow the investigation of a dose-dependent effect of the number of APOE ε4 alleles. Future studies, in larger samples, and with measurements of oxidized LDL could help clarify the effects of the APOE genotype on cholesterol metabolism and the downstream impact on WM microstructure.

### Conclusion

Our findings support the myelin breakdown hypothesis of AD, suggesting that oligodendrocytes’ vulnerability to aging and stressors makes myelin an early target in AD’s pathology [1,9]. Modifiable risk factors for AD (e.g., hypertension, diabetes, dyslipidemia) act as stressors that negatively impact WM health and cognition, especially when combined with familial history and APOE4. We found that WM microstructural changes, especially myelination, were associated with cardiometabolic risk factors in older adults with a family history of AD. Notably, LDL-cholesterol adversely affected WM microstructure only in APOE4 carriers. Our results also suggest that these WM alterations lead to impaired cognition, particularly short-term memory, in APOE4 carriers. This aligns with the theory that genetic and environmental risk factors exacerbate myelin breakdown and accelerate cognitive decline [1,9,79].

## Abbreviations

AD: Alzheimer’s disease
APOE: Apolipoprotein E gene
BMI: Body mass index
BSR: Bootstrap ratio
CSD: Constrained spherical deconvolution
D2: Mahalanobis distance
DTI: Diffusion tensor imaging
DWI: Diffusion-weighted imaging
FA: Fractional anisotropy
FOD: Fibre orientation distribution
GM: Grey matter
HbA1c: Glycated haemoglobin
HDL: High-density lipoprotein
LDL: Low-density lipoproteins
NODDI: Neurite orientation dispersion and density imaging
PLS: Partial least squares analysis
RBANS: Repeatable Battery for the Assessment of Neuropsychological Status
SBP: Systolic blood pressure
WM: White matter

## Author Contributions

Stefanie A Tremblay: Writing - Original Draft, Writing - Review & Editing, Formal analysis, Conceptualization, Methodology, Software, Visualization

Nathan Spreng: Funding acquisition, Resources, Writing - Review & Editing

Alfie Wearn: Formal analysis, Visualization, Writing - Review & Editing

Zaki Alasmar: Methodology, Software, Writing - Review & Editing, Conceptualization

Amir Pirhadi: Software, Methodology

Christine L. Tardif: Writing - Review & Editing, Methodology

Mallar M. Chakravarty: Methodology, Investigation

Sylvia Villeneuve: Funding acquisition, Investigation, Resources, Writing - Review & Editing

Ilana R. Leppert: Software, Methodology, Investigation

Felix Carbonell: Methodology

Yasser Iturria-Medina: Methodology, Writing - Review & Editing, Conceptualization, Funding acquisition

Christopher J Steele: Methodology, Conceptualization, Writing - Review & Editing, Software, Funding acquisition

Claudine J Gauthier: Supervision, Conceptualization, Writing - Review & Editing, Funding acquisition

## Acknowledgments

Data were provided by the PRe-symptomatic EValuation of Experimental or Novel Treatments for Alzheimer’s Disease (PREVENT-AD) program data release 6.0. PREVENT-AD was launched in 2011 as a $13.5 million, 7-year public-private partnership using funds provided by McGill University, the Fonds de Recherche du Québec – Santé (FRQ-S), an unrestricted research grant from Pfizer Canada, the Levesque Foundation, the Douglas Hospital Research Centre and Foundation, the Government of Canada, the Canada Fund for Innovation, the Canadian Institutes of Health Research, the Alzheimer Society of Canada, and the Alzheimer Association. Private sector contributions are facilitated by the Development Office of the McGill University Faculty of Medicine and by the Douglas Hospital Research Centre Foundation (http://www.douglas.qc.ca/).

## Acknowledgments

We would like to acknowledge the PREVENT-AD research group who collected this dataset. The Founders of the program were John C. S. Breitner, MD, MPH, Judes Poirier, PhD and Pierre Etienne, MD, Douglas Hospital Research Centre and Faculty of Medicine, McGill University, Montréal, QC, Canada. Program Current Director is Sylvia Villeneuve, PhD, the Co-Director is Judes Poirier, PhD and the Study Coordinator is Jennifer Tremblay-Mercier, MSc. PREVENT-AD is the result of efforts of many other co-investigators from a range of academic institutions and private corporations, as well as an extraordinarily dedicated and talented clinical and technical assistant staff, students, and post-doctoral fellows. Christine L. Tardif, Ilana R. Leppert, Sylvia Villeneuve, and R. Nathan Spreng are authors affiliated with the PREVENT-AD research team. We would also like to thank the study participants who were recruited from the greater Montréal area and more distant locations in Québec. For up-to-date information, see https://prevent-alzheimer.net/?page_id=42&lang=en.

