## Supplementary Material for "Sex and APOE4-specific links between cardiometabolic risk factors and white matter alterations in individuals with a family history of Alzheimer’s disease"

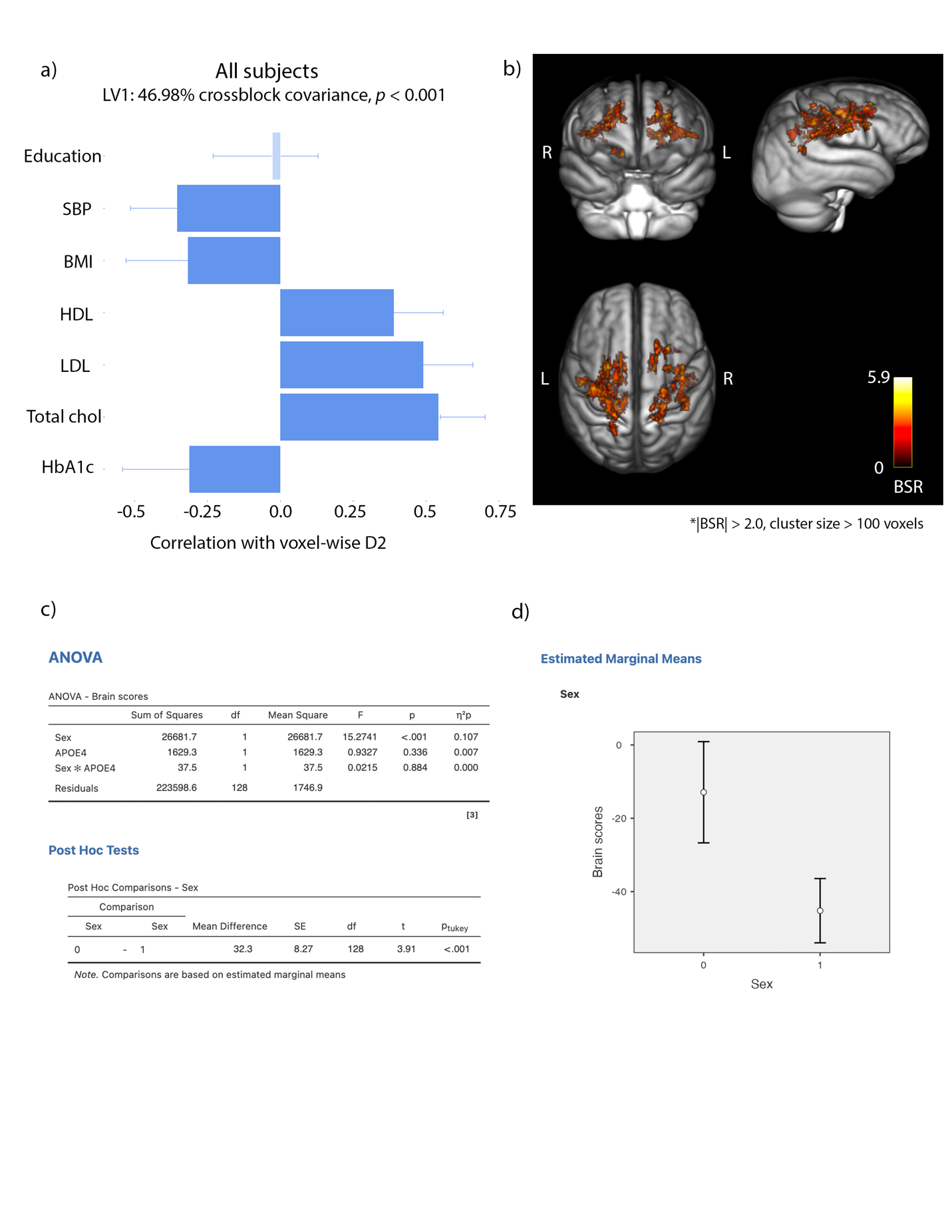

**Supplementary figure 1**. Relationships between D2 in WM and risk factors in the whole sample (males and females). a) The strength and direction of the relationship that each risk factor has with D2 in the voxels shown on the brain images on the right. Error bars show 95% confidence intervals. Correlations are non-significant when confidence intervals overlap with zero. b) Colored voxels (|BSR| > 2.0) have a positive relationship with the patterns shown in (a). The BSR maps are overlaid on a MPRAGE T1w group average image. Higher D2 in the WM regions shown on the right was associated with lower SBP, BMI and HbA1c and with higher HDL, LDL and total cholesterol.

**Supplementary table 1**. Results of the 2x2 ANOVA with sex and APOE4 groups as fixed factors and the brain scores (usc) as the dependent variable.

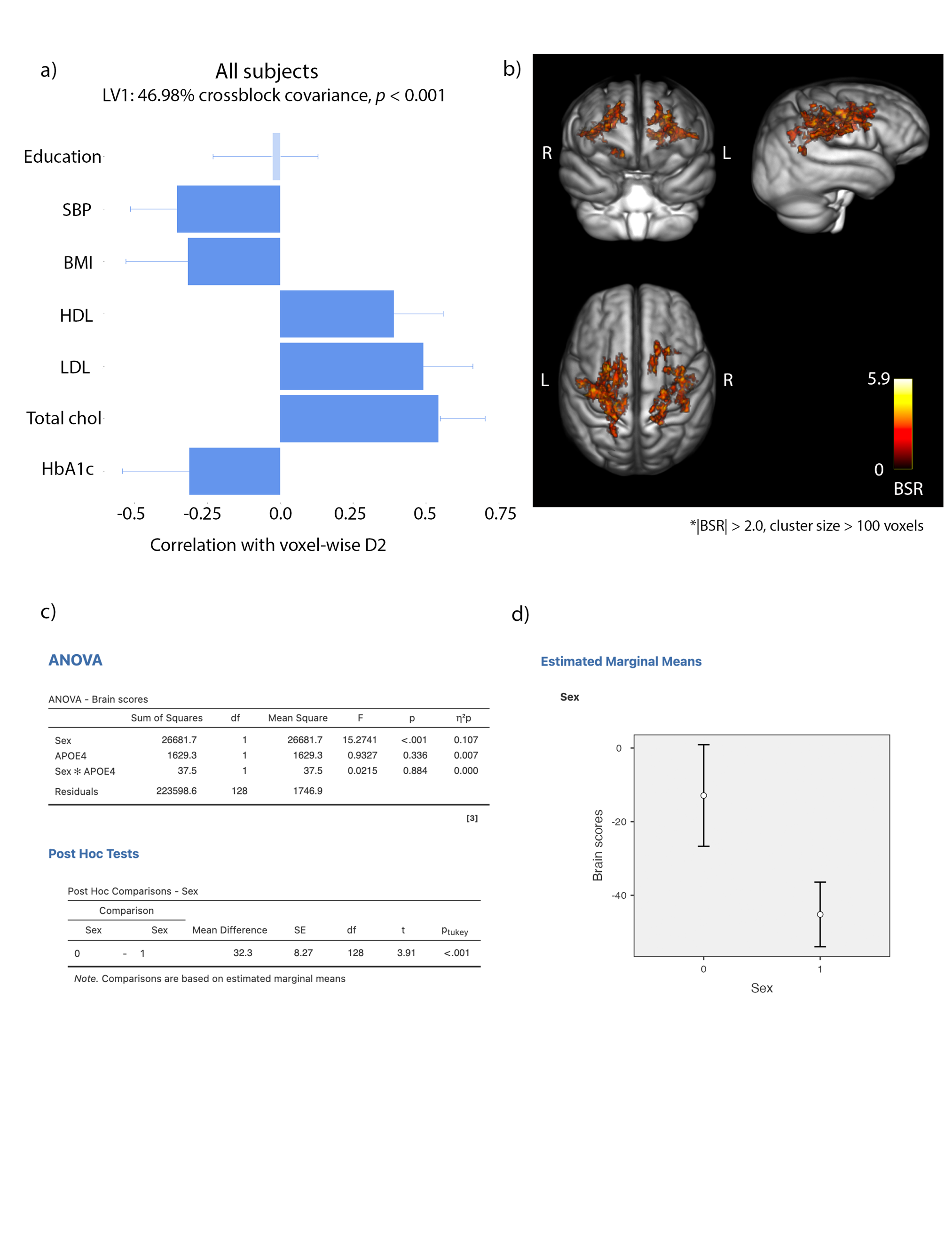

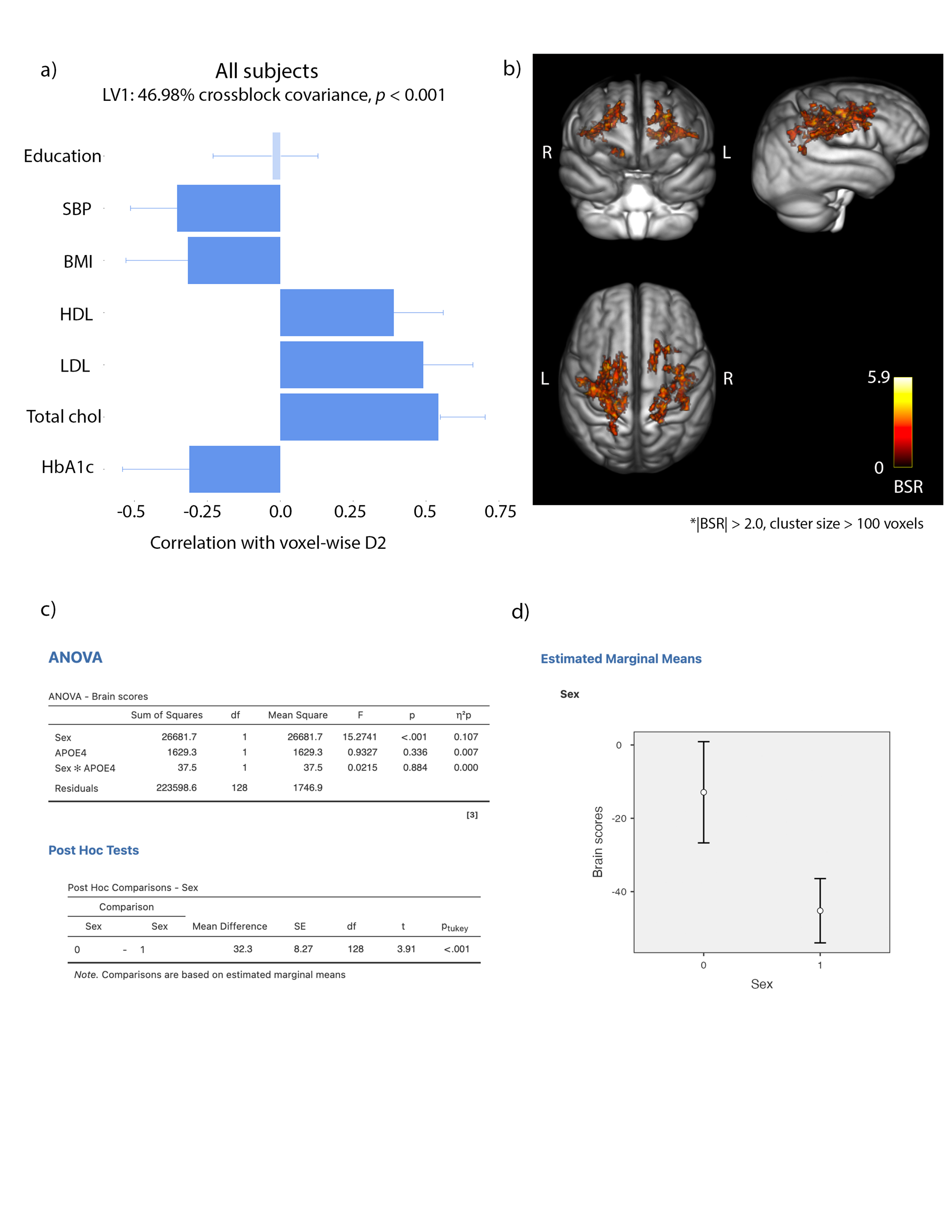

**Supplementary figure 2**. Marginal means plot shows males expressed the pattern of the LV more strongly than females (errors bars = confidence intervals). Males = 0, Females = 1.
